# *Escherichia coli* β-clamp slows down DNA polymerase I dependent nick translation while accelerating ligation

**DOI:** 10.1101/256537

**Authors:** Amit Bhardwaj, Debarghya Ghose, Krishan Gopal Thakur, Dipak Dutta

## Abstract

The nick translation property of DNA polymerase I (Pol I) ensures the maturation of Okazaki fragments by removing primer RNAs and facilitating ligation. However, prolonged nick translation traversing downstream DNA is an energy wasting futile process, as Pol I simultaneously polymerizes and depolymerizes at the nick sites utilizing energy-rich dNTPs. Using an *in vitro* assay system, we demonstrate that the β-clamp of the *Escherichia coli* replisome strongly inhibits nick translation on the DNA substrate. To do so, β-clamp inhibits the strand displacement activity of Pol I by interfering with the interaction between the finger subdomain of Pol I and the downstream primer-template junction. Conversely, β-clamp stimulates the 5’ exonuclease property of Pol I to cleave single nucleotides or shorter oligonucleotide flaps. This single nucleotide flap removal at high frequency increases the probability of ligation between the upstream and downstream DNA strands at an early phase, terminating nick translation. Besides β-clamp-mediated ligation helps DNA ligase to seal the nick promptly during the maturation of Okazaki fragments.

## Introduction

In replicating *E. coli* cells, DNA polymerase III (Pol III) holoenzyme is the major replicative DNA polymerase [1]. Three Pol III core complexes (each containing αεθ polypeptides) interact with the clamp loader complex (γ_1_τ_2_δδ’χψ polypeptides) and β-sliding clamps at the primer-template junctions of a replication fork [2-8]. The dimeric β-clamp is a doughnut-shaped molecule that accommodates double-stranded (ds) DNA at the central hole [2]. When β is free in solution, ATP-bound form of clamp loader complex drives β-clamp opening and binding to the single-stranded (ss) DNA region of a primer-template junction [9-12]. Such a clamp loader interaction with the DNA triggers ATP hydrolysis and results in β-clamp loading on the primer-template junction. Subsequently, the clamp loader affinity to the β-clamp and DNA decreases rapidly that allows clamp loader to dissociate [13,14]. Immediately after being loaded onto the DNA, the sliding clamp interacts with Pol III core at the replication fork [15-17]. The β-clamp also has affinity for all other DNA polymerases in *E. coli* [18-25]. These interactions dramatically increase the processivity of the DNA polymerases [18-25]. Furthermore, interactions of β-clamp with DNA ligase and MutS repair proteins have also been reported [19]. Therefore, apart from acting as a processivity factor for the replicating DNA polymerases, the β-clamp may have some other putative roles in the post-replication DNA repair processes.

Synthesis of the leading strand DNA presumably requires a few priming events [26-30], while frequent priming events at the lagging strand generate discrete Okazaki fragments of 1-2kb in length [31,32]. Maturation of the Okazaki fragments eventually produces an intact lagging strand primarily by the nick translating DNA polymerase I (Pol I) and ligase functions. Pol I is a multifunctional protein with a large and a small domains connected by a flexible linker [33,34]. During nick translation, the large domain of Pol I (also known as Klenow fragment) polymerizes from the 3’ end at the nick and simultaneously displaces 5’ end nucleotides from the template to create variable lengths of oligonucleotide flaps at the nick site [33-37]. Concomitantly, the small domain of Pol I excises the flap by its flap endonuclease activity [33-37]. The strand displacement and flap endonuclease activities are together responsible for the 5’ to 3’ exonuclease (hereinafter 5’ exonuclease) function of Pol I. The Klenow fragment of Pol I also exhibits 3’to 5’ exonuclease (hereinafter 3’ exonuclease) function that is responsible for the proofreading activity [33].

After synthesizing an Okazaki fragment, the replisome dissociates from the β-clamp. As a result, the “left over” β-clamp remains attached at the 3’-side of the ssDNA gap or nick [38-41]. Consistently, β-clamp has been shown to dynamically accumulate behind progressing replication fork in *E. coli* and *Bacillus subtilis* [42,43]. Presumably, β-clamp subsequently helps in repairing the gap or nick by interacting with Pol I, and DNA ligase [19,44,45]. Although a growing body of *in vitro* studies in the past have been done to illuminate the different functions of Pol I, as described above [33-37,46], none of them addressed the dynamics and biochemistry of these functions in the presence of β-clamp. Here, we designed *in vitro* experiments to assess the influence of β-clamp on the functions of Pol I and DNA ligase in the nick translation process. Our approach unraveled a hitherto unknown mechanism of clamp-assisted DNA gap repair in *E. coli.*

## Results

### β-clamp inhibits Pol I-mediated nick translation

To probe the effects of β-clamp on nick translation process, we assembled a template annealing three different oligonucleotides (67, 19 and 28 bases long), as described in Materials and Methods section. Thus, as depicted in the Fig 1A, we created a template that has a primed template junction with a 3’ recessed end of 19 bases long primer and 20 nucleotides single-stranded gap, which suitably acts as a “placeholder” to loaded-β blocking its free sliding [12]. The efficiency and accuracy of primer annealing were almost 100%, as judged by native PAGE (S1 Fig). We loaded β-clamp on the template with the help of clamp loader and ATP, and initiated the polymerization reaction by adding Pol I and dNTPs. The pattern of radioactive oligonucleotide bands in the denaturing gel revealed that Pol I filled the 20 nucleotides DNA gap by extending the radiolabeled 19 bases upstream oligonucleotide and encountered the 28 bases downstream RNA oligonucleotide to initiate nick translation (Fig 1B). Interestingly, β-clamp slightly stimulated pausing of the nick translating Pol I scattered between 39^th^ to 67^th^ positions while traversing 28 bases downstream RNA oligonucleotide (compare the dotted lines in lane 2 and lane 6 of Fig 1B), indicating that β-clamp could moderately reduce the speed of nick translation. On calculating the rate of nick translation, we show that the effect of β-clamp on the Pol I dependent nick translation traversing RNA substrate was nominal (S2 Fig). However, when we used the template with the 28 bases downstream DNA oligonucleotide, the β-clamp dramatically increased the pause of nick translating Pol I, as judged by the bulk migration of the nick sites at 30 and 60 seconds time span (Fig 1C; zones II and I).

**Fig 1.**
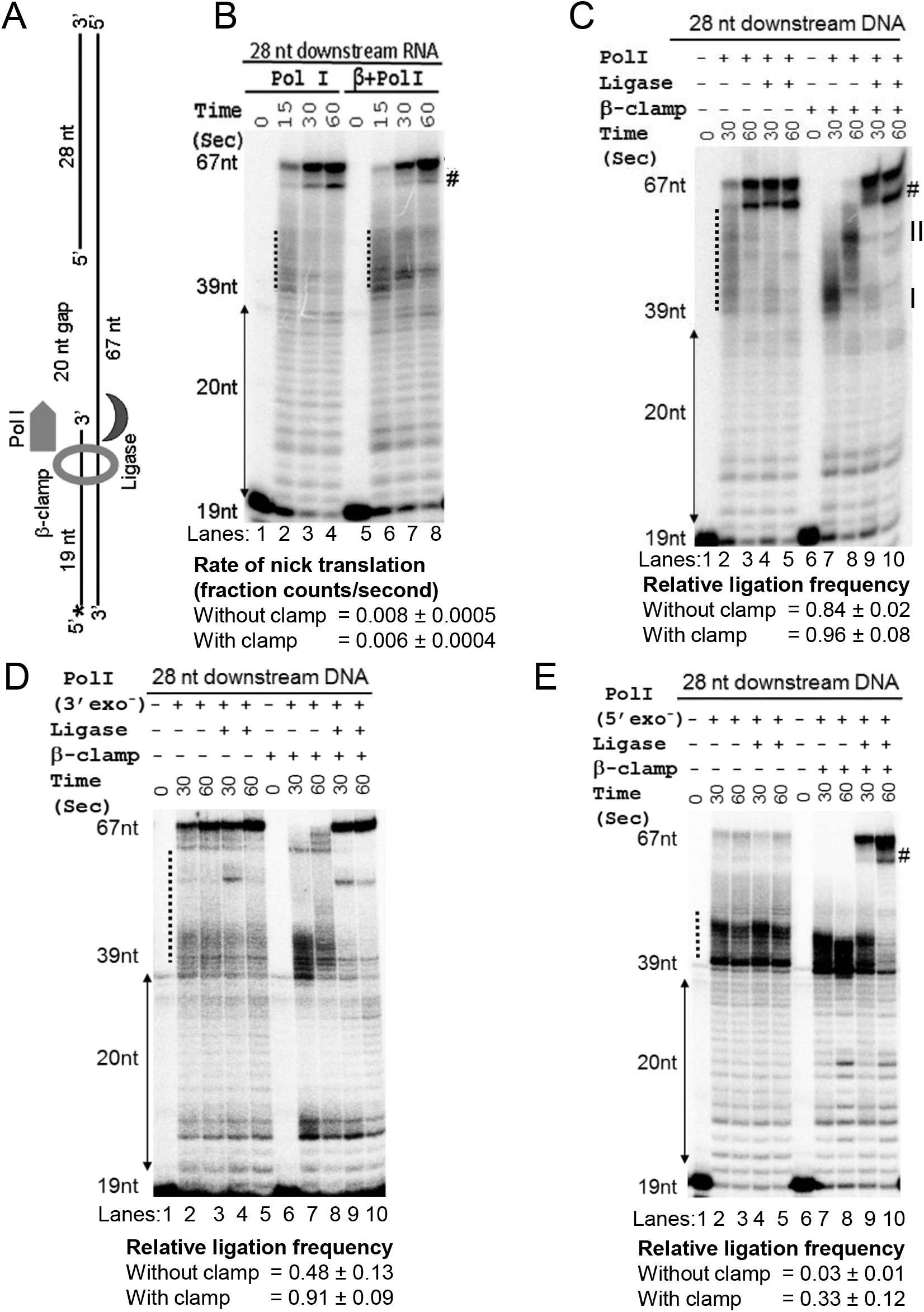
β-clamp inhibits Pol I-mediated nick translation. (**A**) The template used in the assay was prepared by assembling 67 bases, 5’-phosphorylated 28 bases and 5’-radiolabelled (asterisk) 19 bases oligonucleotides. (**B**) A representative urea denaturing PAGE and the rate calculation therein (see also S2 Fig) indicate that the presence of β-clamp nominally affected nick translation traversing 28 bases downstream RNA. (**C**) The urea-denaturing gel shows that Pol I efficiently translated the nick, degrading 28 bases downstream DNA, but the presence of β-clamp dramatically reduced the speed of nick translation. The presence of ligase allowed early ligation when β-clamp slowed down Pol I-mediated nick translation. (**D**) The urea PAGE represents that the 3’-exo^-^ Pol I-mediated nick translation was slowed down in the presence of template-loaded β-clamp, in a manner similar to the nick translation with β-clamp and Pol I combinations, as shown in panel C. Moreover, the presence of template-loaded β-clamp increases the ligation efficiency during the 3’-exo^-^ Pol I-mediated nick translation. (E) The urea denaturing gel represents that the 5’-exo^-^ Pol I intensely paused at the 39^th^ nucleotide and downstream positions, affecting the ligation process. The presence of template-loaded β-clamp moderately enhances these pauses, but facilitates ligation to some extent. The # indicates the position of truncated by-products that originated from the 67-nucleotide long product by the 3’ exonuclease function of Pol I and 5’-exo^-^ Pol I (panel B, C, E). This truncated product is missing in the assay with 3’-exo^-^ Pol I (panel D). All the experiments were performed at least three times. The values represent mean ± standard deviation (sd).

We conducted the control experiments to show that the clamp loader could efficiently load β-clamp on the above-mentioned specific template types (see the detail in S1 text). For this, we used biotinylated templates or histidine-tag version of β-clamp to show that after a clamp loading reaction, the template or the β-clamp could pull-down β-clamp or template, respectively, with 100% occupancy, while clamp loader proteins were washed away (S3 Fig). This observation suggests that the clamp loading was efficient that allows a stable β-clamp interaction with the DNA containing the primer-template junction. Further, to show that the observed effect β-clamp on nick translation is not an artifact arisen by an interference of clamp loader proteins, we performed clamp-loading reaction and washed the streptavidin-bound biotinylated template to remove the clamp loader proteins. Next, we added Pol I and dNTPs to show that the template-bound β could inhibit nick translation similar to the unwashed control reaction containing clamp loader proteins (S4Fig). This data suggests that the template-loaded β-clamp, but not the free clamp loader proteins, in the reaction mixture interfere with the nick translation by Pol I. These observations are consistent with the original literature, which have demonstrated that β-clamp does not slide freely from a template that has a primer-template junction, and the affinity of clamp loader proteins to β-clamp declines rapidly after latter is loaded on the template [9,47].

We incorporated point mutations in *polA* (the Pol I encoding gene) by site directed mutagenesis to disrupt the 3’ and 5’ exonuclease activities. The purified 3’-exonuclease-free (3’-exo^-^) and 5’-exonuclease-free (5’-exo^-^) Pol I were then used to dissect the role of 3’ and 5’ exonuclease activities of Pol I in the nick translation process. The nick translation mediated by the 3’-exo^-^ Pol I (Fig 1D) appeared to be faster than the Pol I-mediated nick translation (compare the intensity of paused bands and 67-nucleotide product between lane 2 of Fig 1C and Fig 1D respectively). To further elaborate this observation, in a separate experiment, we determined that the rate of nick translation mediated by 3’ exo-Pol is faster than the Pol I-mediated nick translation (S5 Fig). However, in the presence of the template-loaded β-clamp, 3’-exo^-^ Pol I-mediated nick translation (Fig 1D) was found to be broadly similar to the Pol I-mediated nick translation patterns (compare lanes 7 and 8 between Fig 1C and Fig 1D). This observation indicates that the 3’-exonuclease activity of Pol I has little impact on the speed of nick translation in the presence of template-loaded β-clamp. On the other hand, the 5’-exo^-^ Pol I exhibited intense pauses at the 39^th^ nucleotide and few bases downstream positions (dotted zone at lanes 2 and 3; Fig 1E). The template-loaded β-clamp modestly enhances the intensity of 5’-exo^-^ Pol I pauses (dotted zone at lanes 7 and 8; Fig 1E). These observations indicate that the elimination of 3’-exonuclease and 5’-exonuclease activities impair the coordination in the strand displacement synthesis process, leading to increased and decreased speed of nick translation, respectively, by Pol I.

### β-clamp inhibits strand displacement function of Pol I

The Klenow fragment of Pol I exhibits polymerization, 3’exonuclease and 5’ to 3’ strand displacement activity [33,34]. To determine if the slow nick translation by clamp-bound Pol I through the DNA substrate was the result of impaired strand displacement, we replaced *E. coli* Pol I with the exonuclease-free Klenow (exo^-^ Klenow), which lacks both 3’ and 5’ exonuclease functions, in the assay. The exo^-^ Klenow exhibited a trail of scattered pause complexes, as judged by the band intensities between the 39^th^ to 67^th^ nucleotide positions (dotted zone at lanes 2 and 3; Fig 2A). Remarkably, the presence of template-loaded β-clamp further intensified the pausing of exo^-^ Klenow, mostly at the 39^th^ nucleotide position (zone III; Fig 2A) where the nick translation begins.

**Fig 2.**
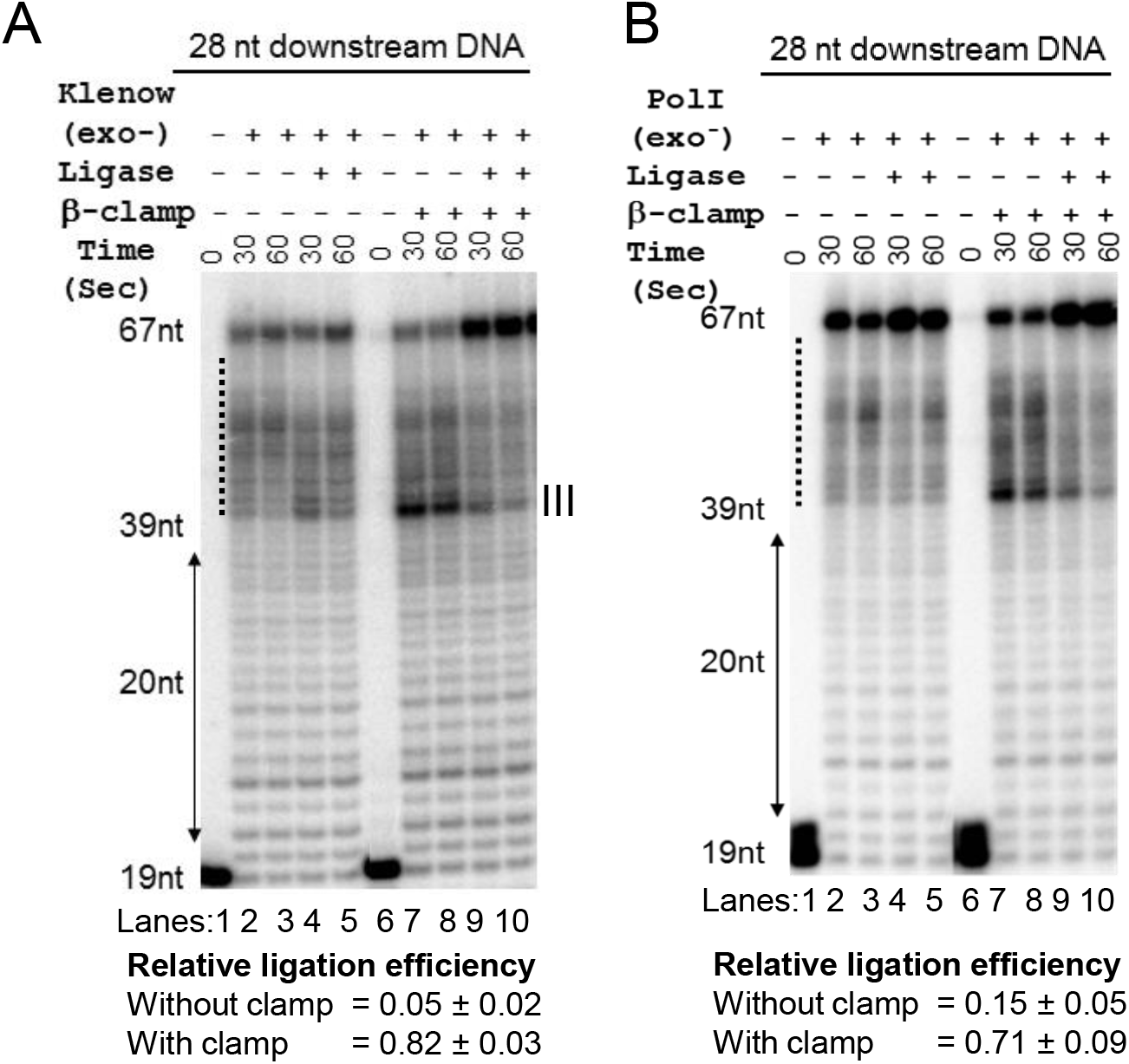
β-clamp inhibits strand displacement activity of exo^-^ Klenow and exo^-^ Pol I. (**A**) The urea PAGE shows that exo^-^ Klenow exhibits a slower speed of strand displacement. Presence of the template-loaded β-clamp stalls the exo^-^ Klenow pauses mostly at the 39^th^ nucleotide position. The exo^-^ Klenow-mediated strand displacement coupled with ligation represents that the DNA ligase fails to ligate nicks. However, the presence of template-loaded β-clamp increases the ligation frequency. (**B**) exo^-^ Pol I exhibits similar functional consequences on the strand displacement and ligation as observed in case of exo^-^ Klenow (panel A). The values representing mean ± sd are calculated from three different experiments.

The choice of exo^-^ Klenow of Pol I, in which the entire small domain containing 5’-exonuclease activity is missing could be non-ideal, because the observed effects might be due to a conformational change rather than stemming from the loss of 5’-exonuclease activity. To test this issue, we generated a mutated version of Pol I, which encodes 3’ - and 5’-exonuclease-free Pol I (exo^-^ Pol I) but retains both of the domains. Using the exo^-^ Pol I, we show that the pattern of paused complexes (dotted zones at the lanes 2 and 3; Fig 2B) was comparable to the paused complexes observed with the exo^-^ Klenow (Fig 2A) in both the absence and presence of the template loaded β-clamp. This data indicates that the elimination of 3’ and 5’-exonuclease activities, but not the deletion of small domain, induce the observed paused complexes in the absence or presence of the template loaded β-clamp. As visually observed, exo^-^ pol I was apparently more efficient in producing 67 nucleotides long product than exo^-^ Klenow in the absence of template loaded β-clamp (Fig 2A and 2B).

### β-clamp promotes ligation at an early stage of nick-translation

Apart from Pol I, DNA ligase also interacts with the β-clamp [19]. Therefore, the polymerization and flap endonuclease activities of Pol I, which fills the ssDNA gap to produce a nick and continuously generates new 5’ ends during nick translation, respectively, may facilitate ligation between the upstream and downstream DNA strands. To probe this working hypothesis, we performed an assay coupled with *E. coli* DNA ligase. We observed that ligase along with Pol I, both in the presence or absence of the template-loaded β-clamp, produced more intense 67 bases long final product than Pol I alone produced in 30 seconds of reaction span (Fig 1C). This observation indicates that ligation by the ligase action during ongoing nick translation quickly contributes creating a fraction of the 67 bases long products. Therefore, almost all of the scattered nick translation products between the 39^th^ and 67^th^ nucleotide positions (marked by the dotted line at lane 2 of Fig 1C), i.e. which were in midway of nick translation, participated in the ligation process to generate 67 bases long product in 30 seconds (lane 4 of Fig 1C). We roughly estimated this contribution of ligase function during nick translation, introducing a new parameter, we termed as “relative ligation frequency” (RLF), that has maximum and minimum values 1 and 0, respectively, by definition (see the Methods section for detail). Thus, RLF is a normalized value or factor that accounts the intensity difference of the 67 bases long product originated in the presence and absence of ligase. Using the formula, RLF is estimated to be 0.84 and 0.96 in the absence and the presence of β-clamp, respectively (Fig 1C) with a statistically insignificant p-value (0.07, T-test) suggesting that β-clamp did not influence the RLF. However, unlike the absence of β-clamp, where the nick translation products were widely scattered (dotted line at lane 2 of Fig 1C), the template-loaded β-clamp restricted nick translation only a few nucleotide downstream of the starting 39^th^ nucleotide position in 30 seconds time span (zone I of lane 7 of Fig 1C). Therefore, the ligase apparently worked at that early nick-point to generate 67 nucleotides long product (lane 9 of Fig 1C). Thus, we infer that by slowing down nick translation, β-clamp also favors ligation at an early stage of nick translation.

We also performed assays to evaluate the RLF while 3’-exo^-^ and 5’-exo^-^ Pol I enzymes take part in the nick translation. The RLF was determined to be less when 3’-exo^-^ Pol I was used in place of Pol I for nick translation assay (Fig 1C and  1D). This data indicates that a faster nick translation by the 3’-exo^-^ Pol I than the Pol I (S5 Fig) inhibits ligase action during nick translation. However, the presence of template-loaded β-clamp, which substantially reduced the speed of 3’-exo^-^ Pol I (Fig 1D), increased the RLF to an extent comparable to the values observed in the assay done with template-loaded β-clamp and Pol I (Fig 1C). In contrast, using the 5’-exo^-^Pol I in the assay blocked the ligation process. This data suggests that in the absence of the 5’-exonuclease function, the 3’-exonuclease and polymerizing centers of Pol I could become hyperactive and thereby blocks access of ligase to the 3’ end of DNA. However, the presence of template-loaded β-clamp with the 5’-exo^-^-Pol I facilitated the ligation incrementally with the duration of reactions (lanes 9 and 10, Fig 1E). These observations indicate that, β-clamp presumably slows down passage through the nick and stimulate 3’-exonuclease function to backtrack Pol I at the nick site facilitating ligation process.

Further, to confirm that β-clamp promotes quick ligation at an early stage, we assessed the influence of β-clamp on the efficiency of ligation when exo^-^ Klenow was employed for the gap filling and strand displacement processes (Fig 2A). The absence of β-clamp severely impaired ligation, as the RLF was determined to be very small (Fig 2A). This data suggests that the strand displacement activity of exo^-^ Klenow pushed the 5’ end of downstream DNA away from the polymerizing 3’ end of the upstream DNA strand (lane 2 of Fig 2A), rendering ligation almost impossible (lane 4 of Fig 2A). On the contrary, the presence of template-loaded β-clamp greatly facilitated the ligation (Fig 2A). The ligation efficiencies were similar when full-length exo^-^ Pol I was used instead of exo^-^ Klenow, both in the absence and presence of β-clamp (compare Fig 2A and 2B).These data suggest that by inhibiting the strand displacement activity, β-clamp extends the duration of the pause of exo^-^ Klenow or exo^-^ Pol I at the 39^th^ nucleotides nick position (lanes 7 and 8 of Fig 2A and 2B). Thus, the β-clamp-mediated weak strand displacement activity of Pol I facilitates DNA ligase to seal the nick efficiently at an early stage (lane 9 of Fig 2A and 2B).

To visually distinguish between the nick translation and nick translation-coupled ligation products, we prepared a modified template replacing 28 bases oligonucleotide with a 38 bases oligonucleotide. Thus, the newly assembled template was identical to the template shown in Fig 1A except for a 10 nucleotide 3’-overhang (S6A Fig). Therefore in a nick translation coupled ligation reaction the full-length product would be 67 bases long if nick translation works alone; while coupled ligation would generate a 77 nucleotide long product. The data presented further emboldened our observation regarding the effect of β-clamp on ligation. The experiments performed with this template and its limitations are elaborated in the S1 text.

In a control experiment, we conducted nick translation-coupled ligation experiment tethering the biotinylated template with the streptavidin beads. After clamp loading, the streptavidin beads were washed to remove the free clamp and clamp loader proteins, as mentioned above. Next, we performed the reaction adding 3’-exo-Pol I, ligase, NAD+ and dNTPs to show only the template-loaded β-clamp, but not free β-clamp or clamp loader complex β-clamp in our above assays, could modulate the ligation function (S7 Fig).

## Loss of DNA contact with the clamp-bound Pol I explains slow nick translation

The finger domain of Pol I interacts with the downstream duplex DNA of the nick site and thereby takes part in displacing the nucleotides of the downstream strand as an oligonucleotide flap [35]. We have depicted this interaction in Fig 3A constructing a model based on the available structures (see Materials and Methods section). Therefore, mutations in the conserved residues of this domain, mainly F771 residue (Fig 3A), inactivate the strand displacement activity of Klenow [30] in a manner similar to the strand displacement activity of β-clamp-bound exo^-^ Klenow and exo^-^ Pol I (Fig 2). Therefore, we hypothesized that β-clamp could allosterically modulate the ternary complex to prevent the interaction between the Klenow and the 5’ nucleotides of the downstream DNA strand. To probe this possibility, we slightly modified the template depicted in Fig 1A, for the assay. Here, we radiolabeled the 28 bases long downstream oligonucleotide instead of the 19 bases long upstream oligonucleotide. Additionally, the 28 bases long oligonucleotide had a bromodeoxyuridine (BrdU) base at the 2^nd^ position. BrdU in the DNA can exhibit a photochemical cross-linking at near UV light on encountering the adjacent amino acids of a binding protein, as described [46]. Using this template, we performed the exo^-^ Klenow-mediated strand displacement assay under UV light. In the absence of β-clamp, a small fraction of BrdU containing oligonucleotide formed the cross-linked adduct with exo^-^ Klenow during strand displacement reaction (lane 2 of Fig 3B), suggesting that the exo^-^ Klenow makes contact with the 28 bases oligonucleotide during strand displacement activity. However, the presence of template-loaded β-clamp abolished the cross-linked adduct, suggesting that β-clamp blocked the contact between the exo^-^ Klenow and the 28 bases oligonucleotide (lane 6 of Fig 3B). Free β-clamp and ATP did not affect the formation of the cross-linked adduct (lanes 3 and 4 of Fig 3B, respectively), suggesting that only template-loaded β-clamp has the said effect. Therefore, the absence of the cross-linked adducts in the presence of β-clamp indicates that the latter affects the interaction between the exo^-^ Klenow and the downstream DNA (Fig 3A), affecting strand displacement function during nick translation (Fig 2).

**Fig 3.**
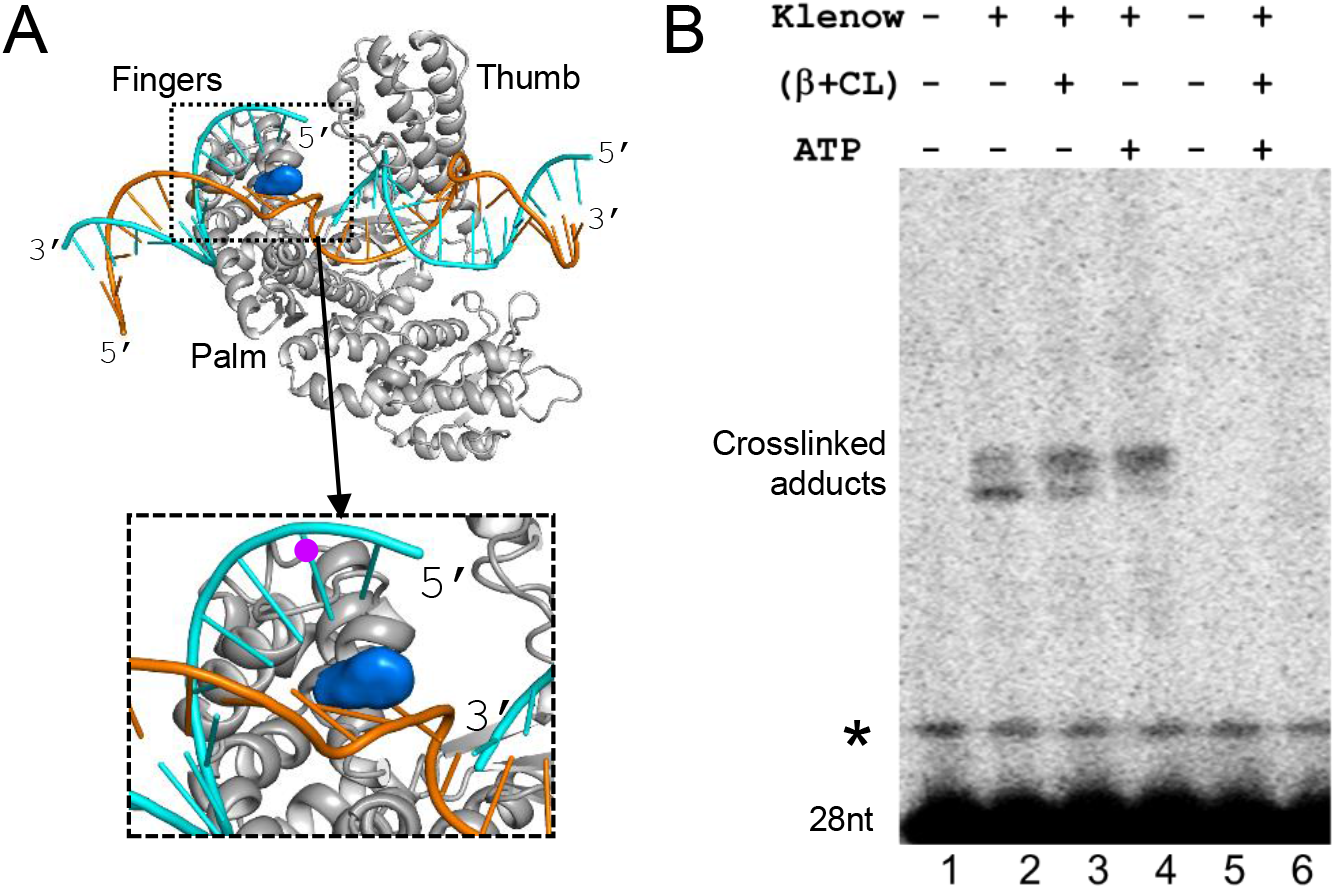
β-clamp inhibits the contact between the 5’end of nicked DNA and Pol-I. (**A**) Structural model of *E. coli* Klenow /DNA complex shows that finger domain of Klenow makes a contact with downstream nick site. The model shows that the conserved F771 (in blue surface representation), which participates in the strand displacement, is positioned between the downstream nicked DNA strands. (**B**) A fraction of BrdU base containing radiolabelled 28 bases oligonucleotide shows a gel shift under the near UV light, suggesting that the oligonucleotide makes cross-linked adduct with Klenow during strand displacement (lane 2). The presence of β-clamp and γ-complex in the absence of ATP (lane 3), or ATP alone (lane 4) also exhibited similar crosslinking adduct. However, β-clamp, γ-complex, and ATP together suppressed the formation of crosslinking product (lane 6), suggesting that loading of the β-clamp on the template blocks the crosslinking. * Represents a minor contaminated band with the custom synthesized 28 bases oligonucleotide.

Moreover, it is noteworthy that unlike static interaction studies [46], the cross-linking assay was essentially performed here in a dynamic reaction (strand displacement) process. As a result, BrdU and the adjacent aminoacyl residue(s) of the finger subdomain of Pol I made a contact in a short time frame (in seconds) under UV light, leading to a poor level of crosslinked adducts (Fig 3B).

### β-clamp stimulates 5’ exonuclease activity of Pol I

The flaps of various lengths generated transiently by Pol I are eventually excised by the flap endonuclease activity of the small domain of Pol I, releasing 1-12nt long products [37]. We show that the flap endonuclease activity of small domain of Pol I in the presence of template-loaded-β-clamp is a crucial step to determine the efficiency of DNA ligase function (Fig 2). Further, to check the effect of template-loaded β-clamp on the flap endonuclease activity of Pol I, we prepared a template where the 5’ end of the 28 bases downstream oligonucleotide was radiolabeled (Fig 4A). We observe that the β-clamp modestly increased the rate of mononucleotide release from the 5’ end of the 28 bases oligonucleotide by Pol I (Fig 4C and S8A Fig). In another case, we used a 5’-end radiolabeled 38 bases downstream oligonucleotide, of which the first 10 bases formed a flap structure at the 5’ end (Fig 4B).

The template-loaded β-clamp enhanced the rate of cleavage of the flap by Pol I to 2.5 folds (Fig 4D and S8B Fig). These data suggest that the template-loaded β-clamp greatly stimulates the flap endonuclease activity of Pol I. We hypothesize that this increased 5’ exonuclease activity of the clamp-bound Pol I for mononucleotides or shorter DNA flaps frequently generate 5’ ends that helps in early and quick ligation, as shown in Fig 1C.

**Fig 4.**
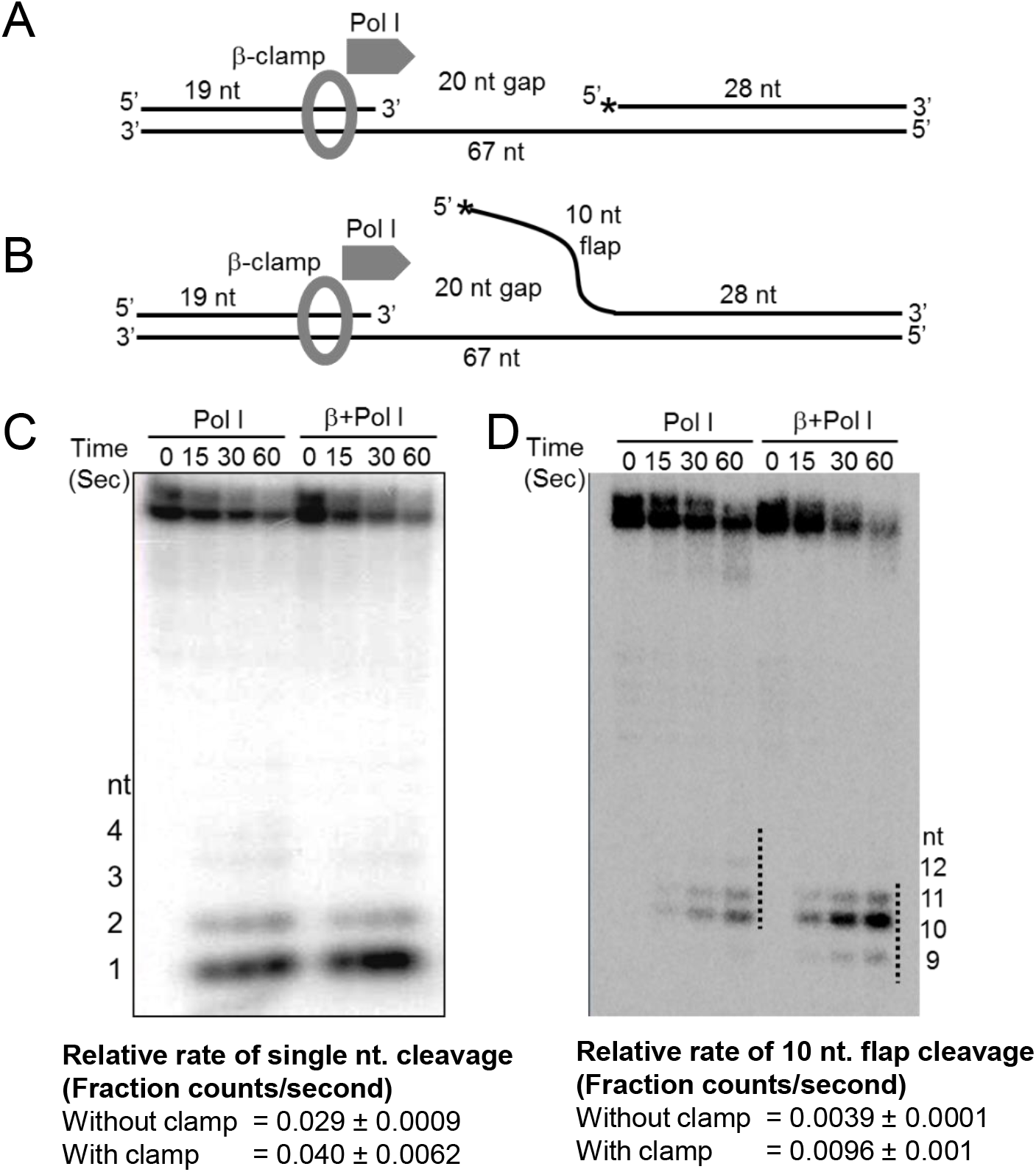
β-clamp increases the flap endonuclease activity of Pol I. **(A)** and **(B)** show the templates assembled using the various lengths of oligonucleotides. Asterisks (*) indicate the radiolabelled 5’-end of the oligonucleotides. **(C)** A representative denaturing gel indicates that the presence of β-clamp modestly increased the efficiency of single nucleotide cleavage at different time points. Also see S8A Fig for the fraction cleavage rate calculation. **(D)** A representative denaturing gel shows that β-clamp greatly enhanced the efficiency of 10 nucleotide long flap removal. Also see S8B Fig for fraction cleavage rate calculation. All the experiments were done at least three times. The values represent mean ± sd.

## Discussion

In brief, we demonstrate that the presence of the template-loaded β-clamp inhibits contact between the downstream strand and the exo^-^ Klenow (Fig 3), impairing the active strand displacement activity of exo^-^ Klenow (Fig 2A). This loss of contact ultimately slows down nick translation by Pol I traversing downstream DNA (Fig 1C). Conversely, β-clamp barely affects the rate of nick translation traversing RNA substrate (Fig 1B and S2 Fig). Since DNA:RNA hybrids are thermodynamically less stable than DNA:DNA hybrids [48,49], we propose that a spontaneous bubbling at the downstream edge of a nick would be greater in the DNA:RNA hybrid. This phenomenon would allow displacement of the strand to be less dependent on the active participation of finger domain, making nick translation easier at DNA:RNA hybrids in the presence of β-clamp (Fig 1B). Such spontaneous bubbling at the ends of DNA:DNA would be lesser due to its higher thermodynamic stability, causing the nick translation to slow down (Fig 1C). Thus, we hypothesize that β-clamp prevents faster nick translation traversing DNA, but not at the RNA primer zone of an Okazaki fragment.

Furthermore, we show that β-clamp enhances the 5’ exonuclease activity of Pol I (Fig 4). To elucidate the possible mechanism by which β-clamp influences the 5’ exonuclease activity of Pol I, we first performed molecular docking between β-clamp and Klenow structures. The model displaying the highest stability under molecular dynamics simulation was chosen (S9 Fig). In addition, the modeling of Klenow/β-clamp/DNA ternary complex supports the feasibility of this Klenow/β-clamp interaction as DNA can follow an overall straight path with minimal strain (S9 Fig). Next, we modeled *E. coli* small domain and attached it to the N-terminal portion of Klenow with a flexible linker to get a final model (S10 Fig). We inferred from the model that the interaction between β-clamp and Pol I occludes the steric conformational space of small domain and pushes it more towards the flap DNA sites. This hypothesis explains the increased flap endonuclease activity of Pol I bound to β-clamp (Figs 3C and D).

Our study also unravels how β-clamp helps in the ligation process to repair nicked DNA. First, β-clamp increases the ligation efficiency by interacting directly with the ligase (Fig 1C) [19]. Secondly, by slowing down nick translation (Fig 1C) and increasing the 5’ exonuclease activity of Pol I (Fig 4), β-clamp promotes early ligation (Figs 1C and 2). The RLF was dramatically reduced when we used 3’-exo^-^ Pol I, 5’-exo^-^ Pol I, exo^-^ Pol I, or exo^-^ Klenow versions of Pol I (Figs. 1D, 1E, and 2), suggesting that the loss of concerted functions of exonucleases and polymerization affected the Okazaki fragment maturation. Interestingly, β-clamp was able to slow down strand displacement activities of the mutated polymerases, thereby restoring the ligation function to different extents. Thus, β-clamp serves as the common interacting platform for replication and repair proteins to maintain overall DNA metabolism and genome integrity by avoiding prolonged polymerization and depolymerization steps, and quickly terminating nick translation, as depicted in the schematic diagram (Fig 5). Therefore, these hitherto unknown aspects of β-clamp function, along with its previously known role as a processivity factor for Pol I and other DNA polymerases [18-25] could have immense evolutionary significance in the efficient maintenance of genome integrity of prokaryotic organisms under challenging environmental milieu.

**Fig 5.**
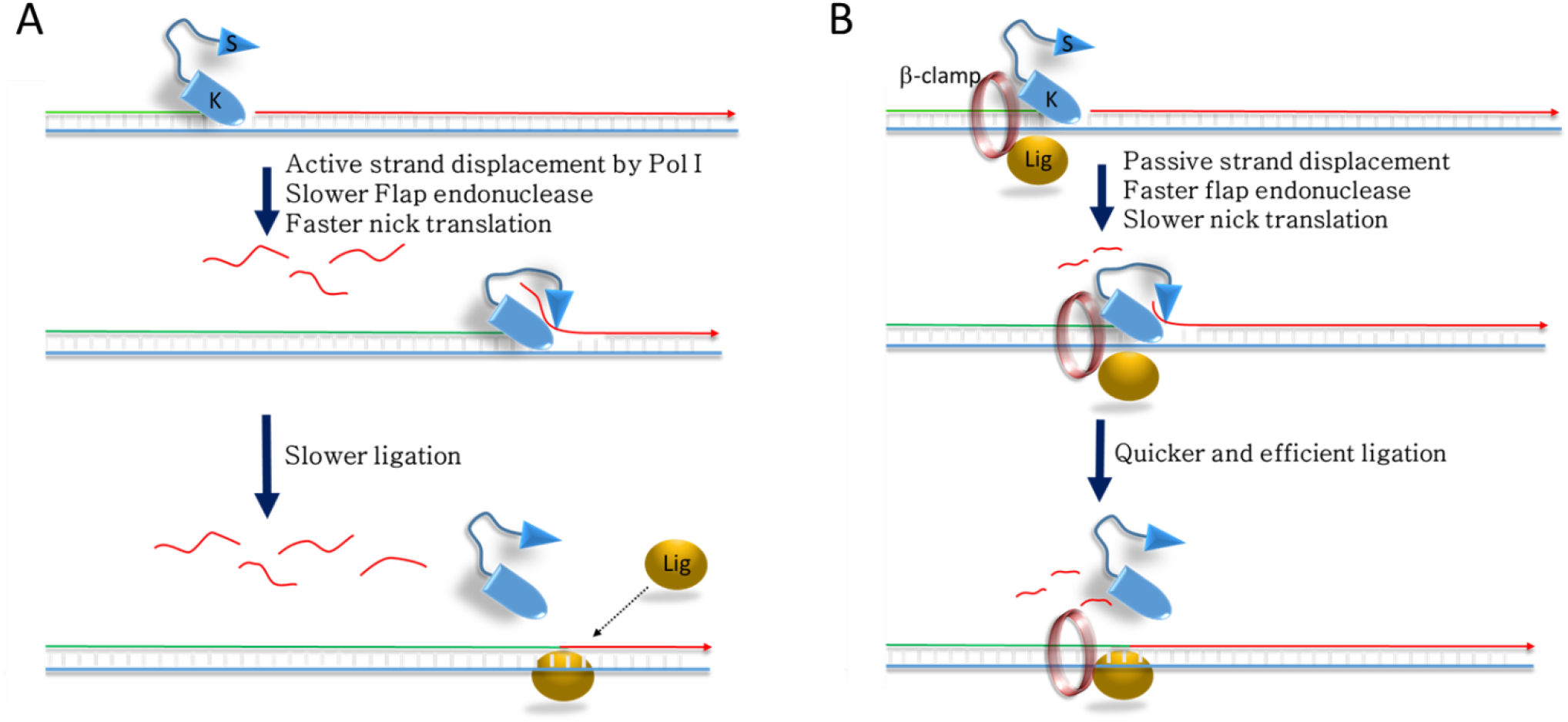
The model showing influence of β-clamp in gap repair. **(A)** The model shows that Pol I removes downstream oligonucleotide by an active nick translation. (red flap).The upcoming ligase molecule may ligate the polymerizing upstream (green) and the depolymerising downstream (red) strands occasionally. **(B)** The clamp bound Pol I has been shown at the nick site. Passive strand displacement for shorter stretch of nucleotides is shown (short red flap). We hypothesize that efficient flap removal of clamp-bound Pol I frequently creates new nick sites. As a result, upcoming ligase molecule may have plenty of time to ligate polymerizing upstream (green) and depolymerising downstream (red) DNA strands at an early stage.

## Materials and Methods

### Proteins and reagents used in the assay

PCR of the genes *(dnaN, dnaX, holA* and *holB)* encoding β-clamp and minimal clamp loader complex (γ,δ and δ’) [50] were performed. Since dnaX gene can produce full length τ or truncated γ subunits, we performed PCR only for the coding region that encompasses γ subunit region [51].The PCR products were cloned at the *NdeI* and EcoRI sites of pET21a (+) vector and expressed as tag-less native proteins. The overexpressed proteins were purified using standard protocols [16,52]. Pol I, exo^-^ Klenow, *E. coli* DNA ligase, and ultrapure nucleotides were purchased from New England Biolabs (NEB). Furthermore, to get full length hexa-histidine-tagged Pol I, PCR amplified *E. coli polA* gene was cloned at the NdeI and HindIII sites of pET28a (+) vector. Site directed mutagenesis was performed on the pET28a-polA vector using Q5 Site-Directed Mutagenesis Kit from NEB to get 3’-exo^-^ Pol I (D^355^A, E^357^A), 5’-exo^-^ Pol I (D^13^N, D^138^N), and exo^-^ Pol I (D^13^N, D^138^N, D^355^A, E^357^A) [45,53]. The hexa-histidine-tagged Pol I and its mutated versions were overexpressed and were purified by nickel-NTA affinity chromatography. The proteins were concentrated, quantified and stored at -20°C in storage buffer (20 mM Tris-HCl, pH 7.5; 500mM NaCl, 1mM DTT and 50% glycerol). Nicotinamide adenine dinucleotide (NAD+) was purchased from Sigma-Aldrich. The hexa-histidine-tagged β-clamp was purified by cloning and overexpressing *dnaN* gene as described in S1 text. The PAGE-purified oligonucleotides (IDT, Inc., USA) used to prepare templates for the assay were, 67 bases (GGATCCCACAC TCATTAAAATTAGTCGCTAATGCATTCTAAAAGCATTCGCAACGAGAAGATAGAGG), 19 bases (CCTCTATCTTCTCGTTGC G), 28 bases (GCGACTAATTTTAATGAGTGTGGGATCC and GCGACUAAUUUUAAUGAGUGUGGGAUCC), and 38 bases (AGAA CTTGACGCGACTAATTTTAATGAGT GTGGGATCC) in lengths. The 5’-end of 28 bases primer was phosphorylated. The biotinylated template made for the control experiments are described in Supporting information.

### Assay conditions

5’-end radiolabeling of the desired oligonucleotides were performed using T4 polynucleotide kinase (NEB) and ^32^P-γATP. The radiolabeled oligonucleotides were purified using BioGel P-6 columns (Bio-Rad, Inc.) to remove the salts and free radioactivity. Templates were prepared as described previously [35,54] using combinations of short oligonucleotides, one of which was 5’-end radiolabeled, as mentioned in the results section. Briefly, 50nM of each oligonucleotide were mixed with 100 μl annealing buffer (20mM Tris-Acetate, pH 7.9, 50mM potassium acetate and 10mM magnesium Acetate). The mixture was heated at 95°C for 2 minutes followed by slow cooling to 37°C to allow annealing of the oligonucleotides into duplex DNA. The formation of duplex was almost complete, as observed in a native PAGE (S1 Fig). To allow clamp loader complex to load β-clamp at the primer template junction, 30 nM β-clamp was incubated with 20nM of annealed template and 30nM clamp loader complex and 1mM ATP in the reaction buffer (20mM Tris-acetate, pH 7.9, 50mM potassium Acetate, 10mM magnesium Acetate, 100μg/ml BSA) at 37°C for 2 minutes. Next, 10μM dNTPs and different versions of Pol I, Klenow and DNA ligase (each at 30nM concentration) were added in the reaction wherever required. For ligation assays, the reaction mixture was also supplemented with 30μM nicotinamide adenine dinucleotide (NAD^+^) to catalyze the ligation process. The final reaction products were separated by urea denaturing PAGE and were exposed to the phosphor-imager screen. A Fujifilm-9000 phosphor-image scanner was used to visualize the bands. Counts of the bands, or the desired stretches of the lanes, were recorded for further analyses. A minimum of three experiments were performed for each reaction to get the average counts and standard deviations. The relative ligation frequency (RLF), i.e. fraction of nick translating products that take part in ligation at 30 seconds time-point, was calculated, as defined below. If, X= average counts of the 67-nucleotide long products generated in the presence of ligase, Y= average counts of the 67-nucleotide long products generated in the absence of ligase, Z = average counts of the products between 39^th^ and 67^th^ nucleotide positions, then RLF = (X-Y)/Z. Nick translation, single nucleotide and 10 nucleotides cleavage rates were calculated from the initial slopes of the fitted curves, as described in the S1 text.

### The BrdU cross-linking assay

The assay was designed with modifications to the protocol mentioned [46]. Briefly, the reaction was performed at 37°C under 310 nm of UV light at 2 cm distance for indicated times to allow cross-linking to occur between the bromodeoxyuridine (BrdU) labeled 28 bases oligonucleotide and nick translating Klenow in the presence or absence of β-clamp. Since the nick translation reaction was observed to reach completion within a minute (Fig 1), we allowed crosslinking for one minute. The reaction mixture was loaded on a denaturing gel to visualize the band shift.

### Structural modeling of *E. coli* Klenow/Nicked DNA complex

The structural modeling of Klenow/nicked DNA complex was done based on the crystal structure of *Geobacillus stearothermophilus.* Klenow complexed with 9 base pairs of duplex DNA (PDB ID 1L3S). The additional upstream and downstream DNA segments were built using COOT (See S1 text).

## Acknowledgements

Authors are grateful to Dr. Jayanta Mukherjee, Bose Institute, India, Dr. Gursharan Kaur, IMTECH, India and Dr. James Boroweic, NYUMC, USA, for reading the manuscript and providing critical suggestions on the manuscript.

## Supporting information

### S1 Fig. Gel shift assay shows correct primer annealing during preparation of template

The native gel shows the separation of 19 nucleotides (lane 1), 28 nucleotides and 67 nucleotides (lane 3) long oligonucleotides. When they mixed in an annealing reaction, the individual oligonucleotides disappeared from their original position and gave rise to a higher band (Lane 4), suggesting that annealing was 100% efficient. However, non-complementary oligonucleotides (19 and 28 bases) did not found to be annealed (lane 5). On the other hand, the 67 bases long oligonucleotide, which has complementary regions for both 19 and 28 bases long oligonucleotides, has also completely annealed to these 19 and 28 bases oligonucleotides (lanes 6 and 7, respectively).

### S2 Fig. β-clamp does not affect the rate of nick translation through the RNA substrates

The fraction of the initial counts that appeared at 67 nucleotide position were plotted. The rate of the nick translation through RNA substrate was calculated from the fitted curve shown above (also see S1 text). The negligible difference between the rate of nick translation in the presence and absence of β-clamp, suggests that β-clamp has no influence on nick translation through RNA substrates.

### S3 Fig. Interaction of the loaded-β clamp on the primed template

(A) The lanes 2-5 are showing purified individual proteins that are used in the pull down assay. The streptavidin-bound primed biotinylated template could pull-down β-clamp (marked by white arrow), while clamp loader proteins washed away (compare unwashed and washed lanes 6 and 7, respectively). (B) The his-tag β clamp pull down primed template, as indicated by the pulled down radiolabeled 19nt long oligonucleotide, which was the integral part of the template.

### S4 Fig. Clamp loader proteins in the reaction mixture do not interfere with nick translation

The urea denaturing gel indicating that washing the streptavidin-bound reaction products, which washes out the clamp loader protein, but retains β clamp (S3 Fig), generates similar pauses (+wash lanes) as observed for standard reaction (-wash lanes) performed in the presence of β clamp loader proteins

### S5 Fig. Comparison between Pol I and 3’-exo^-^ Pol I mediated nick translation

(A) The autoradiogram represents nick translation product formation at three different time points by Pol I and 3’-exo^-^ Pol I. (B) The rate of nick translation was calculated as described in S1 text, from three independent autoradiograms and plotted. The initial rate of nick translation was found to be faster in case of 3’-exo^-^ Pol I.

### S6 Fig. Influence of β-clamp on nick translation coupled ligation

(A). A new template was assembled using 67 bases, 5’-phosphorylated 38 bases and 5’-radiolabelled (asterisk) 19 bases oligonucleotides, so that a 10 nucleotide long 3’ overhang was generated as shown. (B) The autoradiogram represents Pol I mediated nick translation-coupled ligation in the presence or absence of β-clamp. Visibly, 77 nucleotide ligation product that appears at lower time points, degraded quickly due to 3’ exonuclease activity of intact Pol I. (C, D) The representative autoradiograms show that 3’-exo^-^ Pol I and exo^-^ Pol I produced stable nick translation-coupled ligation products in the presence or absence of β-clamp in three different time points. (E) RLFs calculated at different time points in the presence or absence of β-clamp from panels S6C and S6D are shown.

### S7 Fig. The template-loaded β-clamp enhances the efficiency of ligation

The nick translation coupled ligation assay was performed on streptavidin beads to remove clamp loader proteins after loading of the β-clamp. Addition of 3’ exo-Pol I and dNTPs exhibited pausing at the 39^th^ and downstream locations at two different time points. Addition of the Pol I and ligase efficiently produces 67nt products, suggesting that in the absence of clamp loader complex the nick translation coupled ligation reactions are similar as demonstrated in Fig 1D.

### S8 Fig. β-clamp increases flap endonuclease activity by Pol I

(A). Single nucleotide cleavage was shown in Fig 4C. The single nucleotide fraction cleavage values at different time points were plotted to calculate the rate. The presence of β-clamp was found to modestly increase the single nucleotide cleavage. (B) Similarly, flap cleavage was shown in Fig 4D and the flap fraction cleavage values at different time points were plotted to calculate the flap fraction cleavage rates. Interestingly, β-clamp greatly enhanced the flap cleavage rate.

### S9 Fig. Model for PolK/ β-clamp/Nicked DNA ternary complex

(A) Cartoon representation of the docked model of Klenow (cyan) /β-clamp (gray and blue) protein complex. The possible protein-protein interaction interfaces are encircled in red and blue. (B) Molecular dynamics simulations for the docked Klenow/β-clamp protein-protein complex. (C) Surface representation of the modelled Klenow (grey) /β-clamp (yellow and blue)/ DNA (orange and green) ternary complex.

### S10 Fig. Final model for PolI/clamp/Nicked DNA complex

The structural model of *E. coli* small domain of Pol I (light pink) was built using Prime module in the Schrodinger software suite using 1BGX as the template. The models were manually placed near the Klenow/nicked DNA (A) and Klenow/ β-clamp/nicked DNA (B) models using PyMOL. The predicted flexible linker region connecting small domain and Klenow fragment has been shown by undulating lines in different grey shades. The conformational space available in β-clamp-bound and β-clamp free Klenow/nicked DNA structures are shown by multiple copies of Psmall domains. From this model, we hypothesize that the presence of β-clamp could sterically restrict (shown by red T blocker) the conformational freedom of small domain hence increasing the chance of accessing 5’ exonuclease site.

### S1 Text.

**Supporting methods and results**

